# Matrix stiffness affects spheroid invasion, collagen remodeling and the effective reach of stress into the ECM

**DOI:** 10.1101/2025.03.14.643241

**Authors:** Klara Beslmüller, Rick Rodrigues de Mercado, Gijsje H. Koenderink, Erik H.J. Danen, Thomas Schmidt

## Abstract

The extracellular matrix (ECM) provides structural support to cells thereby forming a functional tissue. In cancer, the growth of the tumor creates an internal mechanical stress which, together with remodeling activity of tumor cells and fibroblasts, alters the ECM structure leading to an increased stiffness of the pathological ECM. The enhanced ECM stiffness in turn stimulates tumor growth, activates tumor-promoting fibroblasts and tumor cell migration, leading to metastasis and an increased therapy resistance. The connection between internal tumor stress, ECM stiffness, ECM remodeling, and cell migration are unresolved. Here we used 3D ECM-embedded spheroids and hydrogel-particle stress sensors, to quantify and correlate internal tumor-spheroid pressure, ECM stiffness, ECM remodeling, and tumor cell migration. 4T1 breast cancer spheroids and SV80 fibroblast spheroids showed increased invasion - described by area, complexity, number of branches and branch area - in a stiffer, cross-linked ECM. On the other hand, changing the ECM stiffness only minimally changed the radial alignment of fibers but highly changed the amount of fibers.. For both cell types, the pressure measured in spheroids gradually decreased as the distance into the ECM increased. For 4T1 spheroids, increased ECM stiffness resulted in a further reach of mechanical stress into the ECM which, together with the invasive phenotype, was reduced by inhibition of ROCK-mediated contractility. By contrast, such correlation between ECM stiffness and stress-reach was not observed for SV80 spheroids. Our findings connect ECM stiffness with tumor invasion, ECM remodeling, and the reach of tumor-induced mechanical stress into the ECM. Such mechanical connections between tumor and ECM are expected to drive early steps in cancer metastasis.

## Introduction

Metastasis is the primary cause of cancer related mortality, accounting for over 90% of cancer-related deaths [1]. Metastatic cancer is challenging to treat because it can be diffusely localized in various organs and it is often resistant to cytotoxic agents. Understanding the processes driving cancer metastasis, such as dissemination from the primary tumor and invasion may lead to improved treatment and prognosis of cancer patients.

As a tumor grows and expands it applies pressure onto the surrounding tissue. In addition, tumors actively remodel their surrounding tissue, including the extracellular matrix (ECM). The ECM is a network of glycoproteins, including collagen, providing mechanical support as well as signaling cues to cells [2]. Composition, structural organization, and stiffness of the ECM are altered in tumors with increased stromal stiffness being associated with more aggressive cancers [3, 4]. Within the tumor microenvironment (TME) multiple cell types, including tumor cells and fibroblasts, act in concert to stiffen the ECM. Remodeling of fibrilar collagens in the ECM through enhanced deposition, degradation, and cross-linking is an important aspect of ECM stiffenening. Collagen cross-linking by fibroblasts and tumor cells can help tumor progression by promoting focal adhesion formation and increasing integrin binding [5]. Secretion of lysyl oxidases (LOX) and lysyl hydroxilase 2 (LH2) that contributes to cross-linking is increased in cancers. Targeting LOX has been suggested as a therapeutic avenue for cancer patients [6].

Confinement of tumors growing inside a tissue combined with ECM stiffening and application of traction forces drives mechanical stress and pressure buildup in tumors, which in turn can activate mechanosensitive pathways [7]. A stiffer matrix can be pro-tumorigenic, it can activate tumor-promoting fibroblasts, it can trigger chemoresistance, and it has been associated with increased cancer risk in fibrotic organs [8, 9, 10, 11]. While a stiff ECM may be thought of as a barrier for cell migration, ECM stiffness also induces promigratory morphological changes in tumor cells, epithelial-mesenchymal transition (EMT), invasion, metastasis, and resistance to chemotherapy [12, 13, 14, 15]. Indeed, cancer spheroids showed enhanced migration in a stiff ECM as they exert higher integrin-mediated traction forces [16]. ECM fibers create migration tracks that promote cancer cell invasion [17, 18] and radially aligned collagen fibers at the tumor-stroma interface facilitate local invasion [19]. A stiffer ECM was also shown to promote breast cancer progression by promoting the appearance of cancer-associated fibroblasts (CAFs) [20].

Rho-Rho-associated protein kinase (ROCK) signaling drives cell contractility and application of force on collagen matrices [21]. Simultanously, mechanical forces, generated by the cellular cytoskeleton, are required for active cell movement [22]. Vice versa, elevated ECM stiffness activates FAK/RhoA/ROCK and PI3K/AKT signaling pathways via integrins [23]. The stiffness of the ECM has a major influence on the traction forces, as increased substrate stiffness triggers higher cell traction forces [24]. This positive feedback loop of ECM stiffness activating ROCK, and ROCK activation increasing tissue stiffness, is believed to be a key process in promoting the invasiveness of cancer.

Taken together, solid tumors generate pressure in tissue and alter local tissue mechanics by remodeling the ECM. Such tissue stiffening, in turn, activates tumor growth, invasion, and metastasis. Preventing or reversing tissue stiffening in tumors may have potential to reduce cancer progression [25]. Yet, the connection between tumor pressure, ECM stiffness, ECM remodeling, and tumor cell migration are unresolved. Recent technological advances permit the measurement of traction forces even in a complex 3D environment [26, 27]. Here, we use an innovative 3D traction force technique [28], applying soft elastic hydrogel micro-spheres to measure mechanical stress and pressure in and around ECM-embedded spheroids derived from tumor cells versus those derived from fibroblasts. This stress analysis is combined with correlative quantitative measurements of ECM remodeling and cell migration. Our results connect strains and pressures within spheroids and their reach into the surrounding ECM, to the stiffness of the ECM and to cell migration, offering new insights into the mechanobiology of tumor progression.

## Materials and methods

### Microparticle synthesis

Microparticle synthesis and functionalization was performed similar to previously described [28]. In short, deformable acrylamide-co-acrylic-acid microparticles were synthesized using Shirasu porous glass membranes (SPG Technology, Japan). These tubular membranes with tunable pore size enabled us to produce spherical particles in large quantities and of uniform size in a range of 5-50 µm. A solution of 150 mM NaOH, 0.3% (v/v) tetramethylethelenediamine (TEMED; Thermo Fisher Scientific, Waltham, MA, USA, 17919) and 150 mM 3-(N-Morpholino)propanesulfonic acid (MOPS) sodium salt (Sigma-Aldrich, Waltham, MA, USA, M9381) was supplemented with acrylamide (AAm), acrylic acid (AAc) and cross-linker N,N’-methylenebisacrylamide (BIS) (all Sigma-Aldrich, A9099, 147230, 146072, respectively) with a final pH of 7.4. The total mass concentration of acrylic components was *c*_*T*_ = *c*_*AAm*_ + *c*_*AAc*_ + *c*_*BIS*_ = 100 mg/mL. The relative concentration of acrylamide was set to 10%. A cross-linker concentration *c*_*c*_ = *m*_*BIS*_*/*(*m*_*AAm*_ + *m*_*AAc*_ + *m*_*BIS*_) of 1.5% was used. The Young’s modulus of the particles was subsequently characterized using atomic force microscopy. The Young’s modulus was *E* ≈ 600 Pa.

### Microparticle functionalization

First, particles were washed in activation buffer (100 mM MES (Sigma-Aldrich, M5057) and 200 mM NaCl), and subsequently incubated for 15 min in reaction buffer of 0.1% tween20 (Sigma-Aldrich, P7949), 4% (w/v) EDC (Sigma-Aldrich, E7750) and 2% (w/v) NHS (Thermo Fisher Scientific, 22500). Microparticles were washed and incubated for 1 h with 5 µg/mL of BSA in PBS for SV80 experiments, and 5 µg/mL E-cadherin-FC in PBS for 4T1 experiments at pH 8. Subsequently, Alexa647-Cadaverine (Thermo Fisher Scientific, A30679) was added for 30 min. Unreacted NHS groups were blocked using 300 mM Tris and 100 mM ethanolamine (Sigma-Aldrich, 398136) (pH 9). Finally, particles were washed thrice in PBS with 0.1% tween20 and stored for use in PBS (pH 7.4) with 5 mM sodium azide.

### Cell culture

Cell lines SV80 and 4T1 (obtained from the ATCC) were cultured in high-glucose Dulbecco’s Modified Eagle’s Medium (DMEM, Gibco, Thermo Fisher Scientific, Waltham, MA, USA, 11504496) containing L-Glutamine and Sodium Pyruvate, supplemented with 10% fetal calf serum (Thermo Fisher Scientific) and 25 µg/mL penicillin/streptomycin in a humidified incubator at 37°C with 5% CO_2_.

### Rheology

To test the glutaraldehyde (GTA) efficiency in cross-linking collagen, we performed rheology measurements (Anton Paar, Graz, Austria, Physica MCR 501) on collagen gels in cone-plate geometry, between stainless steel 40-mm-diameter plates and 0.992° truncation angle. Collagen type I solutions were isolated from rat-tail collagen by acid extraction as described previously [29]. Collagen matrices were created by mixing with DMEM, HEPES 0.1 M (1 M stock, Sigma-Aldrich, H0887) and NaHCO_3_ 44 mM (Sigma-Aldrich, S8875) to a collagen end concentration of 2 mg/mL as previously described [30]. The plates were heated to 37°C prior to loading the sample. Water was added to the solvent trap as to maintain a moist environment. 28 min after loading the sample, a solution of either pure DMEM or DMEM supplemented with 0.1% GTA (Sigma-Aldrich, 340855) was placed around the geometry and allowed to diffuse into the sample for 1 h. The elastic and viscous moduli of the network were probed every 5 seconds by applying an oscillatory deformation of 0.5% strain amplitude at a frequency of 0.5 Hz. Measurements were recorded and analyzed using software provided by the manufacturer (Anton Paar, RHEOPLUS/32 v3.41D100712).

### Spheroid formation

The collagen and buffers described in the previous paragraph were used for spheroid embedding. Collagen matrices were created by mixing with DMEM, HEPES, NaHCO_3_, and growth factor-reduced basement membrane matrix, matrigel^®^ (Corning, Corning, NY, USA, 354230).

Matrigel was added to further decrease the stiffness of the gel. Final collagen concentrations of 0.6 mg/mL, and matrigel concentrations of 1.5 mg/mL were used. Fluorescent microparticles were added to the collagen mixture to a concentration of 80 particles/µL. 30 µl of the collagen mixture was polymerized in 384-well CELLSTAR^®^ plates (Greiner Bio-one, Kremsmünster, Austria, 781091) at 37°C for 1 h. Non-enzymatic cross-linking was performed by incubating the gel with 0.1% GTA for 1 h. To ensure soluble GTA was completely removed, gels were subsequently washed with PBS 8 times, then twice with medium [31].

Spheroids were injected into the 3D collagen-Matrigel matrix as described previously [32]. In short, subconfluent monolayers of tumor cells were trypsinized and filtered (Sysmex, Norderstedt, Germany, 04-0042-2317). Cells were mixed with fluorescent microparticles (E-cadherin coated were used for 4T1, BSA coated microparticles were used for SV80) in a ratio of approximately 1000:1 and resuspended in PBS containing 2% polyvinylpyrrolidone (PVP, Sigma-Aldrich, P5288) to reach a cell concentration of approximately 50000 cells/µl. Immediately thereafter, spheroids, 200 µm in diameter and containing approximately 2500 cells, were created by automated injection of the cell-PVP mixture into the collagen-matrigel matrix at defined x-y-z positions 150 *µ*m above the bottom of the wells using an injection robot (Life Science Methods, Leiden, The Netherlands). After injection, spheroids were incubated with the appropriate medium, or medium containing 20 nM Rho kinase inhibitor (ROCK1/2 GSK 269962, Tocris, Bristol, UK, 4009) at 37°C. As 4T1 tumor cells grew and migrated faster than SV80 fibroblast cells, 4T1 spheroids were incubated for 48 h, and SV80 spheroids for 72 h. All spheroids were fixed and stained with final concentrations of 2% formaldehyde (Sigma-Aldrich, 252549), 0.1% TritonX-100 (Sigma-Aldrich, T8787), 0.4 µg/mL Hoechst (Fisher Biotech, Wembley, Australia, 33258) and 0.05 µM AlexaFluor-488 Phalloidin (Invitrogen, Waltham, MA, USA, A12379) in PBS for 3 h at room temperature. The samples were washed thoroughly in PBS followed by microscopy.

### Microscopy

#### Scanning confocal microscopy

Images of spheroids were acquired on a Nikon C2plus Ti2 inverted scanning confocal microscope equipped with four laser lines 405/488/561/640 nm, and with a SimpleSI detector. The microscope has a Nikon encoded and automated stage. Its camera is controlled through NIS Element Software (Nikon Instruments Inc., Melville, NY, USA). Spheroids were imaged with 20 µm distance between z-slices. ECM collagen-fibers were detected by confocal reflection-microscopy on a Nikon Eclipse Ti inverted scanning confocal microscope equipped with an A1R MP scanner. The collagen fibers were scanned at 561 nm excitation with a 561 nm blocking dichroic. All light in the range 400 - 750nm was collected on a GaAsP-photomultiplier. Scanning confocal fluorescent microscopy of spheroids, and scanning confocal reflection microscopy of the collagen networks were performed using Plan-Apo ×20/0.75 NA, and Apo-LWD ×20/0.95 objectives (Nikon Instruments Inc.), respectively.

#### Spinning-disk confocal microscopy

High-resolution imaging was performed on an inverted microscope (Zeiss, Oberkochen, Germany, Axiovert 200) equipped with a 20X, 0.5 NA Plan-Neofluar objective (Zeiss). The setup was expanded with a confocal spinning-disk unit (Yokogawa, Musashino, Japan, CSU-X1), and an emCCD camera (Andor Technology, Belfast, Northern Ireland, iXon DU897).The hydrogel microparticles were imaged with 642 nm diode laser illumination (Spectra Physics, Utrecht, Netherlands). A 561 nm DPSS-laser (Cobolt, Stockholm, Sweden) was used to image the AlexaFluor-561 Phalloidin. A 405 nm DPSS-laser (CrystaLaser, Reno, US) was used to illuminate the Hoechst nuclear-stain. Z-stacks were used to obtain a 3D image-cube of the microparticles and the surrounding cells. The mismatch between the refractive indices of the air surrounding of the objective and the culture medium caused an apparent compression of the image cube, which was corrected by an experimentally-determined correction factor: the apparent elongation of at least 10 stiff (> 5 kPa) microparticles was determined by calculating the radius in the equatorial plane compared to the lateral radius of the particles. The ratio of both was used as a correction factor for the z-positions, rendering the effect of the refractive index mismatch negligible.

#### Collagen fiber analysis

To analyze the orientation and alignment of collagen fibers, reflection images were processed using a custom Matlab (R2021a) script. First, at the mid-plane of the spheroid, one z-plane of a reflection image stack was taken to generate a representative image of the collagen fibers. The center of mass (CoM) of the spheroid was determined and the spheroid was separated from the background. The image was blurred with a narrow Gaussian kernel. The orientation and coherency (alignment) of the fibers was quantified using this image [33]. These parameters were assessed at the fiber locations by means of a weighted mean over the signal intensity, and the radial alignment of the fibers was calculated relative to the CoM of the spheroid. The distance of the fibers to the edge of the spheroid was calculated using the spheroid’s foreground image. To facilitate comparison of the profile of radial fiber alignment and coherency of many spheroids, the data was binned with a bin size of 15 µm based on the distance to the edge.

#### Migration analysis

The home-built migration analysis in Matlab was based on a method described previously [34]. Scanning confocal z-stacks of the actin cytoskeleton were projected using the standard deviation in the z-direction (Fig. S1 A). A Gaussian filter with a narrow kernel was used to remove small fluctuations from the projected image. An adaptive threshold was used to separate the foreground from the background. The area *A* and perimeter *P* of the spheroid were calculated using the the generated foreground image. For quantitation we defined a normalized complexity value *C* as

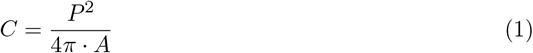

denoting the roughness of the spheroid edges. The point in the foreground that was farthest from the background was defined as the centroid and its distance to the background as the core radius (green dot and circle in Fig. S1 B). The invasive area was defined by the area of the foreground outside the core. To identify branches, skeletonization was applied to the foreground mask, with the endpoints of the resulting skeleton serving as branch points (cyan dots in Fig. S1 B-C).

#### 3D spheroid reconstruction

The center of mass (CoM) and radius of a partially imaged spheroid were determined using software coded in Matlab (Fig. S2). A Gaussian filter with a wide kernel was applied to the image stacks containing the actin cytoskeleton. Subsequently, a 3D Sobel operator was used the highlight the edges of the spheroid. The foreground was separated from the background and a least square non-linear fit was used the fit a sphere to the foreground voxels. The center and radius of this sphere defined the CoM and radius of the spheroid.

#### High-resolution microparticle shape reconstruction

Z-stacks of particles were taken on a spinning-disk confocal fluorescence microscope. The z-slices were separated by at a distance of Δ*z* = 1.2 µm which is half the depth-of-focus of the objective to comply to Nyquist’s theorem. Individual particles in the image-cube were identified, and the data cubes cropped to contain solely individual particles.

#### Local stress analysis

Fast stress analysis was performed in spherical harmonics space [35] implemented in the Python SHTools library [36]. In short, spherical harmonics are solutions to the generalized linear elasticity continuity equation (Hooke’s law) for the mechanical equilibrium condition of a spherical boundary in an isotropic medium. Deformations were expressed by complex coefficients, *û*_*jlm*_, of the spherical harmonics:

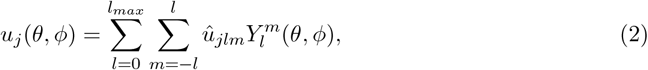

where 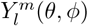 are the set of complex spherical harmonics functions given by the harmonic-degree *l* and order *m*. Using those solutions of the displacement field, the full stress tensor was calculated in terms of it’s complex stress coefficients 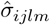 which are uniquely defined by the displacement coefficients 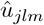:

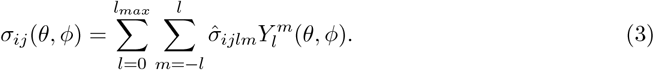

Any local stress on the microparticles leads to a local deformation of the particle’s surface at the location of the stress. We defined indentations as negative stress. The full stress tensor was subsequently separated into an isotropic (pressure, *P*), and an anisotropic (deviatoric stress, D) component:

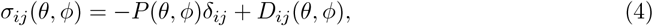

where *δ*_*ij*_ refers to Kronecker delta. Since the deviatoric stress is traceless by definition, the pressure is expressed as:

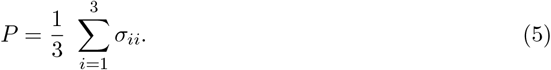

The spatial resolution of the stress field was limited by the maximum harmonic degree, *l*_*max*_, in the spherical harmonics expansion. Here we used *l*_*max*_ = 15, since this gives a good balance between spatial resolution (≈ 2 µm), and computational time (≈ 1 h per particle on an office PC).

#### Statistics

A one-tailed, unpaired, parametric t-test was used with Welch’s correction, in order to determine statistical significance between two one-dimensional populations. For comparison of two-dimensional datasets (i.e. those containing both x- and y-coordinates), a contrast-to-noise ratio (CNR) was calculated:

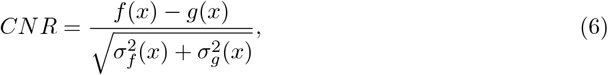

where *f* (*x*) and *g*(*x*) are datasets depending on a spatial variable *x. σ*_*f/g*_ are the respective standard deviations. A one-sample t-test on *CNR* was used to determine statistical significance between *f* (*x*) and *g*(*x*). Data sets were significantly different with probabilities of *p* < 0.05 (*), *p* < 0.01 (**), and not significantly different for probabilities of *p* > 0.05 (*ns*).

## Results

### Glutaraldehyde cross-linking increases collagen network stiffness without affecting collagen fiber architecture

In order to test the effect of glutaraldehyde (GTA) on the elastic properties of collagen matrices, we measured the elastic modulus of collagen by rheology. Collagen is a viscoelastic material for which the mechanical properties are determined by two components. The storage modulus *G*^*′*^, which represents the elastic component of a viscoelastic material, accounting for the amount of energy that is stored in the material during deformation, and the loss modulus *G*″, which represents the viscous component, accounting for the amount of energy that is dissipated during deformation.

*G*′and *G*″ were measured at a frequency of 0.5 Hz during polymerization of 2 mg/ml collagen. The collagen gels polymerized within 10 minutes as displayed in Fig. S4 A. Subsequently, GTA was added while *G*′and *G*″ were further monitored during the process of cross-linking. On GTA cross-linking, a 4.5-fold increase in storage modulus (to 91 ± 7 Pa, mean ± sem) was observed after 1 hour incubation with 0.1% GTA, as compared to incubation with control medium (20 ± 5 Pa) (Fig. 1 A). No change in the loss modulus was observed after adding GTA, as shown in Fig. 1 B.

**Figure 1:**
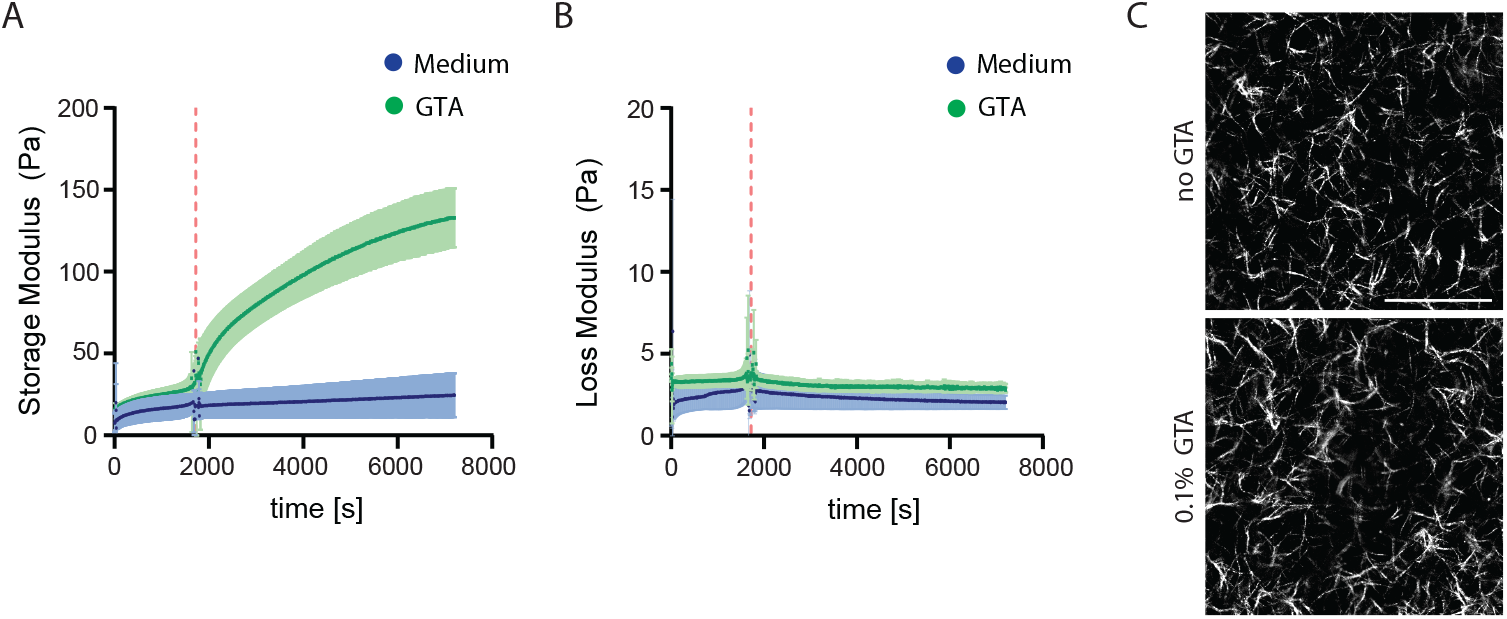
GTA stiffens the ECM but does not change the collagen fiber architecture. **A-B**. Rheology measurements of collagen gels (Storage and Loss modulus respectively). The shaded area shows the standard deviation of the mean. The red dashed line indicates the time point in which either GTA or medium was added to the gel. **C**. Reflection confocal microscopy of collagen gels. Scalebar: 100 µm.

The architecture of the collagen gels with and without GTA was analyzed using reflection microscopy. No difference in fiber length and fiber diameter was observed as shown in Fig. 1 C. We further quantified fiber density in three independent experiments using Fiji (National Institutes of Health, USA, version 1.53c)[37]. A threshold was applied to a maximum projection of 5 z-slices, and the area covered by the fibers was quantified (Fig. S4 B). No significant increase in fiber density in GTA gels was observed as compared to control collagen gels. Likewise, the collagen architecture appeared similar in gels with and without GTA cross-linking.

These results indicated that GTA cross-linked collagen gels behave more elastic, store more energy, and return to their original shape more readily (*G*′ higher). They do not exhibit increased viscous deformation (*G*″ similar), and the collagen fiber architecture is not affected by cross-linking.

### Stiffness increases cell invasion into the ECM for spheroids derived from 4T1 breast cancer cells and for fibroblasts

Spheroids of SV80 fibroblasts and 4T1 breast cancer cells were cultured in GTA-cross-linked and non-cross-linked gels for three and two days, respectively. In pure 2 mg/ml collagen gels GTA had no effect on spheroid growth and morphology. In order to expose the spheroids to even softer environments, where the impact of cross-linking was predicted to be more robust, we mixed 0.5 mg/ml collagen with 1.5 mg/ml matrigel for all further experiments. The inclusion of matrigel did not affect the collagen architecture (Fig. S3). Those gels were softer than *G*′< 10 Pa, the resolution limit of our rheometer. We extrapolated that cross-linking with GTA would likewise lead to 4.5-fold increase in storage modulus as we had observed for 2 mg/ml pure collagen gel. Spheroids were imaged after fixation and staining. Fig. 2 A shows an overview of representative spheroid images.

**Figure 2:**
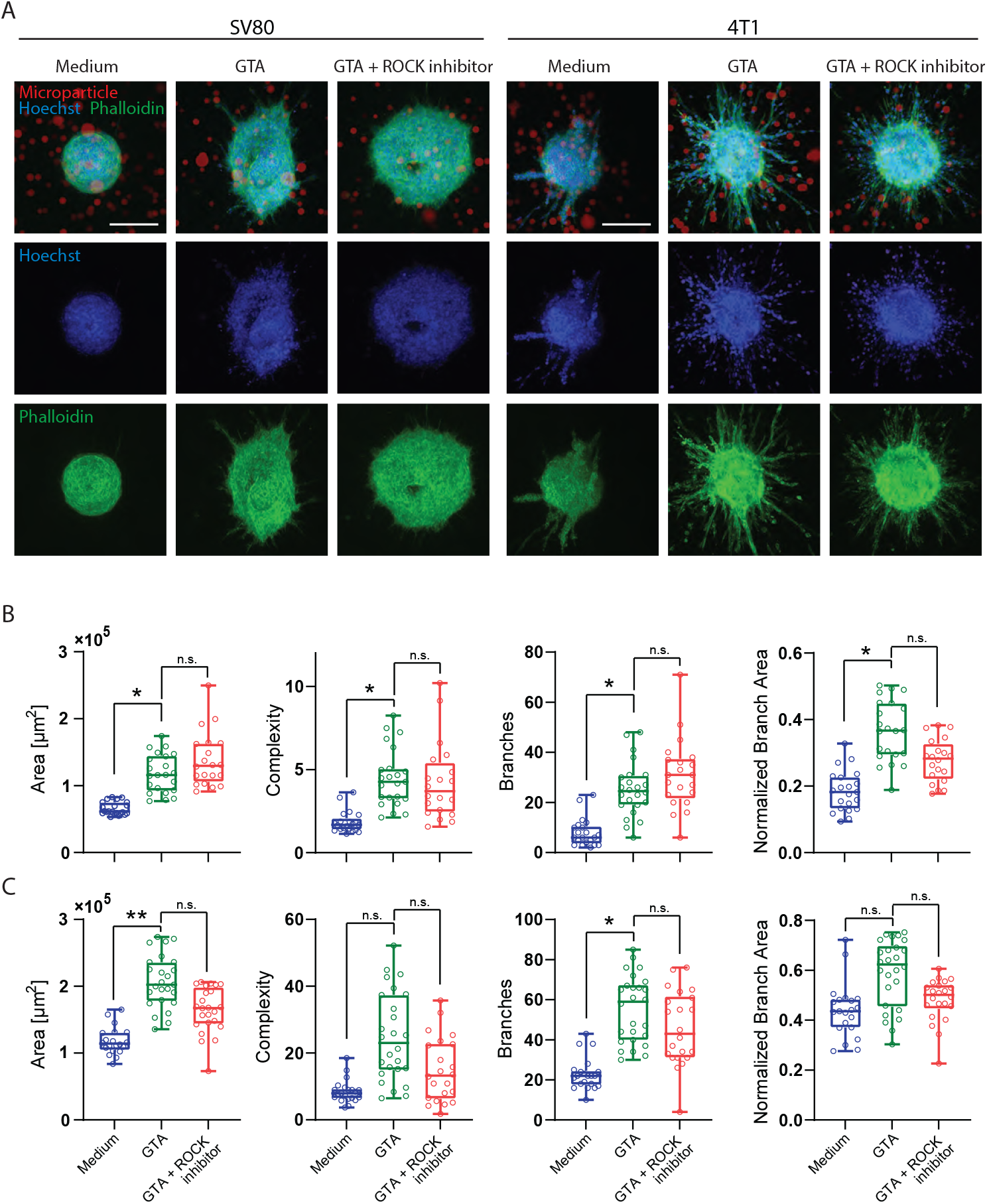
Migration of spheroids. **A**. Maximum projections of SV80 and 4T1 spheroids after 72 and 48 h, respectively, cultured in soft gel (medium) and stiff gel (GTA). Scalebar: 200 µm. **B**. Area, Complexity, Number of branches and branch area analysis of SV80 spheroids from three independent experiments. All spheroids from three experiments are plotted in the graphs. Calculations on p-values are done with n=3. **C**. Area, Complexity, Number of branches and branch area analysis of 4T1 spheroids from three independent experiments. All spheroids from three experiments are plotted in the graphs. Calculations on p-values are done with n=3.

Images were analyzed in the phalloidin channel, labeling the cellular F-actin architecture, using a home-built spheroid invasion analysis tool. Based on a binary foreground separation of the spheroids, spheroid area, complexity, branch number, and branch area were calculated and compared between all conditions. Fig. 2 B-C summarizes these results for SV80 (B) and 4T1 (C) spheroids, respectively. For both cell types, spheroids showed a significant increase in area, complexity as a measure spheroid’s edge ruffling, branch number, and normalized branch area in GTA cross-linked gels as compared to control gels. A minimum of 20 spheroids each was analyzed. The spheroid area was increased by a factor of 1.76 (sem medium: 0.06; sem GTA: 0.10) for 4T1, and 1.80 (sem medium: 0.05; sem GTA: 0.11) for SV80. The complexity was increased by a factor of 2.9 (sem medium: 0.13; sem GTA: 0.4) for 4T1, and 2.4 (sem medium: 0.11; sem GTA: 0.3) for SV80. Notably, while 4T1 and SV80 spheroids showed a similar area, 4T1 displayed significantly higher complexity (≈ 4-fold) and branch area (≈ 2-fold) as compared to SV80 spheroids.

As ROCK-mediated traction forces are known to increase in stiffer ECM, we asked whether the increased migration into cross-linked ECM may be ROCK dependent. For this, 20 nM of the ROCK1/2 inhibitor GSK 269962 was added to spheroids in GTA gels. Indeed, for both 4T1 and SV80 spheroids exposure to ROCK1/2 inhibitor led to a decrease in complexity and branch area parameters (Fig. 2 B-C). However, while the total spheroid area and the number of branches was likewise decreased for 4T1 cells in the presence of GSK 269962, SV80 spheroids showed a slight increase in spheroid area and number of branches upon ROCK1/2 inhibition. Thus, for 4T1 but not for SV80 spheroids, the inhibition of ROCK1/2 led to a shrinkage in cross-linked gels, becoming more similar to behavior in soft gels in absence of the inhibitor.

Togeter, this data demonstrated that stiffening of the ECM increases cell invasion out of spheroids for both cell types. However, the role of ROCK1/2-mediated contractility and traction, was different for the two cell types.

### Cell-type and stiffness affects ECM remodeling

We next analyzed how 4T1 and SV80 spheroids remodeled the surrounding ECM fiber-network, and how remodeling was affected by ECM stiffness. In soft ECM, both 4T1 and SV80 spheroids aligned the fibers of the collagen in radial direction centered to the spheroid core. 4T1 spheroids thereby displayed much higher capacity to pull and create thick collagen bundles as compared to SV80 spheroids (Fig. 3 A). Remarkably, while GTA cross-linking appeared to slightly increase the number of thin radially oriented fibers around SV80 spheroids, the fiber network around 4T1 spheroids was depleted. For both cell types, inhibition of ROCK1/2 minimally affected collagen remodeling, confirming our observations with respect to spheroid invasion above.

**Figure 3:**
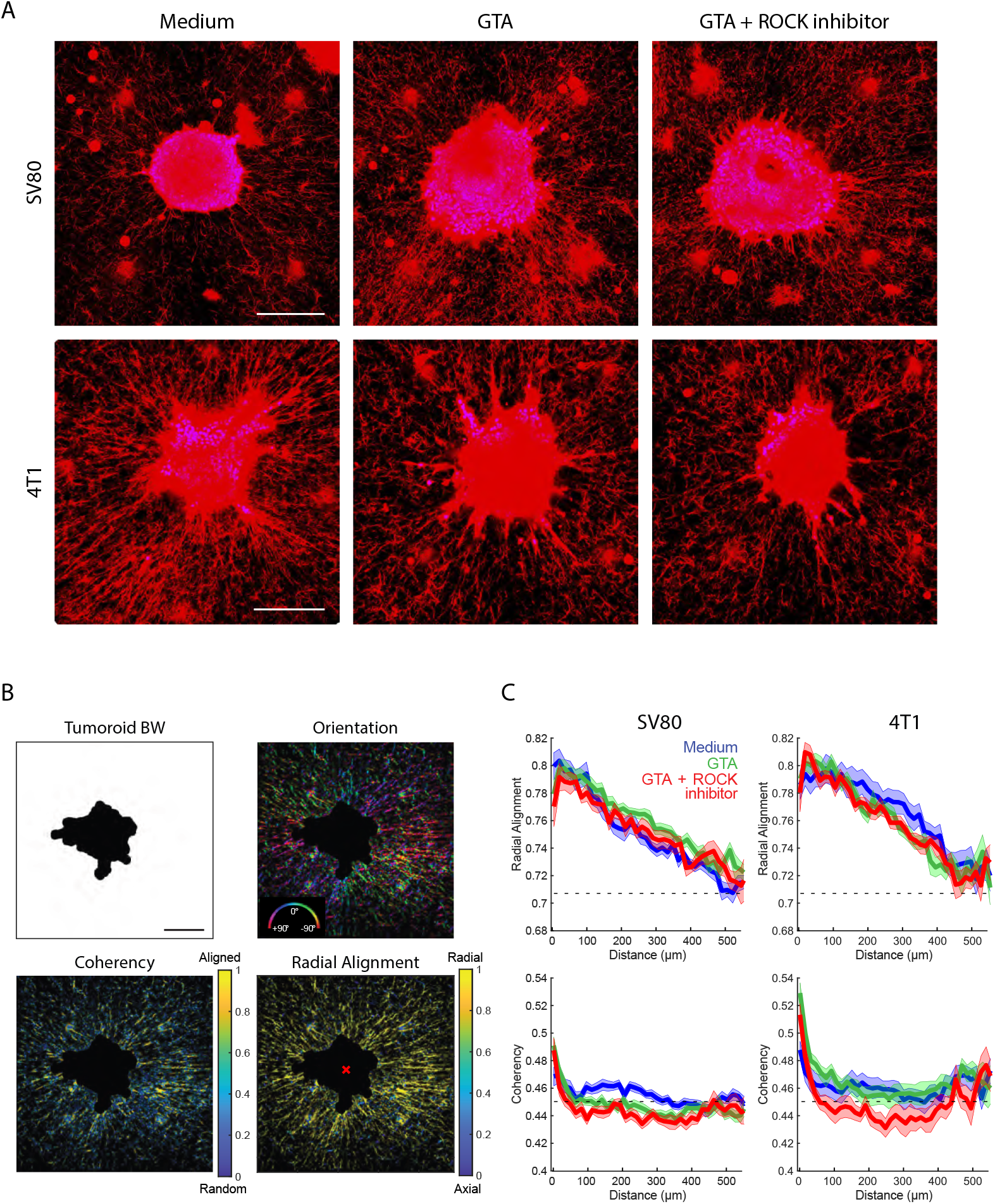
Radial collagen alignment around spheroid is affected by ECM stiffness. **A**. Overview of representative reflection images of collagen fibers for each experimental conditions. Scalebar: 200 µm. **B**. Representative orientation and coherency decomposition of a 4T1 spheroid in medium condition. The distance to the spheroid was calculated using the shortest distance to the spheroid foreground (spheroid BW). Radial alignment was calculated relative to the centroid of the spheroid (red x). Scalebar: 200 µm. **C**. Average radial alignment and coherency for *≥* 12 reflection images of collagen fibers per condition. The shaded area shows the standard error of the mean. Close to the edge of the spheroid, fibers showed high coherency and were aligned in the radial direction. With increasing distance from the spheroid’s edge, the alignment dropped to a random network (dashed line, control) at around 500 µm.

For quantitative analysis of the changes in the organization of the ECM network, we quantified the orientational order of the structure that emerged. The fiber orientation (the direction of fibers relative to the spheroid), the radial alignment relative to the centroid of the spheroid, and the coherency (a length scale of constant fiber alignment) were determined, following the methods developed by Püsköki *et*.*al* [33] (Fig. 3 B). The radial alignment decreased gradually from 0.8 at the spheroid edge, to 0.71 at about 500 µm from the spheroid edge. The latter value is predicted for a random network. The overall behavior was similar for all spheroids and for all conditions (Fig. 3 C). In agreement with the visual results shown in Fig. 3 A, the reach of ECM remodeling into the collagen gel was longer for 4T1 spheroids as compared to that for SV80 spheroids. The typical length scales found were 400 and 250 *µ*m, for 4T1 and SV80, respectively.

The coherency was increased especially close to the spheroid’s edge, which may point to active mechanical stresses applied by the cells that increase the length scale at which fibers remain aligned (Fig. 3 C). Notably, the coherency measures the alignment without a given direction, which results in a non-zero offset for a random network. The coherency decreased rapidly from SV80 and 4T1 spheroid edges into the ECM approaching the inherent fiber coherency value of 0.45 within 100 µm from the spheroid edge. Following this initial drop, coherency remained slightly above background and reached the inherent fiber coherency after 300 µm for both cell lines. In a stiffer gel, coherency at the spheroid border was higher for both cell lines. Moreover, it was maintained further into the surrounding ECM, in a ROCK-dependent manner, for 4T1 but not SV80.

These findings showed that spheroids derived from both cell types strongly reorganize the collagen network on length scales of hundreds of micrometers, and that ECM stiffness has a limited impact on organization or length scale.

### Matrix stiffness affects reach of spheroid-induced stress into the ECM

The stress originating from spheroid expansion and the traction forces applied by the cells of the spheroid onto the ECM will extend into the ECM as inferred from the emergence of fiber orientational order shown above. We quantified the local mechanical stress and pressure distribution inside and around spheroids using soft elastic hydrogel microparticles. Microparticles were mixed into the ECM solution before polymerization, and mixed with the cells before injection into the ECM. We noticed that uncoated microparticles were rapidly excluded from the spheroids. Therefore, we coated the microparticles with the collagen-binding protein E-cadherin prior to co-injection into the preformed gel for 4T1 spheroids. We observed that such coated microparticles readily attached to the 4T1 cells and remained within the spheroids. As E-cadherin expression was absent in SV80 cells (Fig. S5) we used BSA-coated microparticles for SV80 spheroids, which likewise prevented exclusion of the microparticles from SV80 spheroids.

The local pressure fields were determined from the microparticle deformations as detailed in the M&M section to an accuracy of ≈ 40 Pa (Fig. 4 A-B). As we simultaneously tracked the absolute position of each microparticle with respect to the spheroid centroid we were able to reconstruct the local stress and pressure fields throughout the sample. One of the critical observations was the increase in absolute pressure (more negative pressure/indentation, i.e. compression) of the microparticles towards the core of the spheroid as shown in Fig. 4 C. The pressures leveled off with distance from the spheroid core. Our data revealed that pressure extended beyond the spheroid edge (the zero on the x-axis of the graphs) into the surrounding ECM. Microparticles far off from any spheroid (*r* > 500 µm) reported on pressures of 40 Pa (dashed lines in the graphs) set by the accuracy of our measurements. The results for each individual biological replicate is shown in Fig. S6 of the supplementary material.

**Figure 4:**
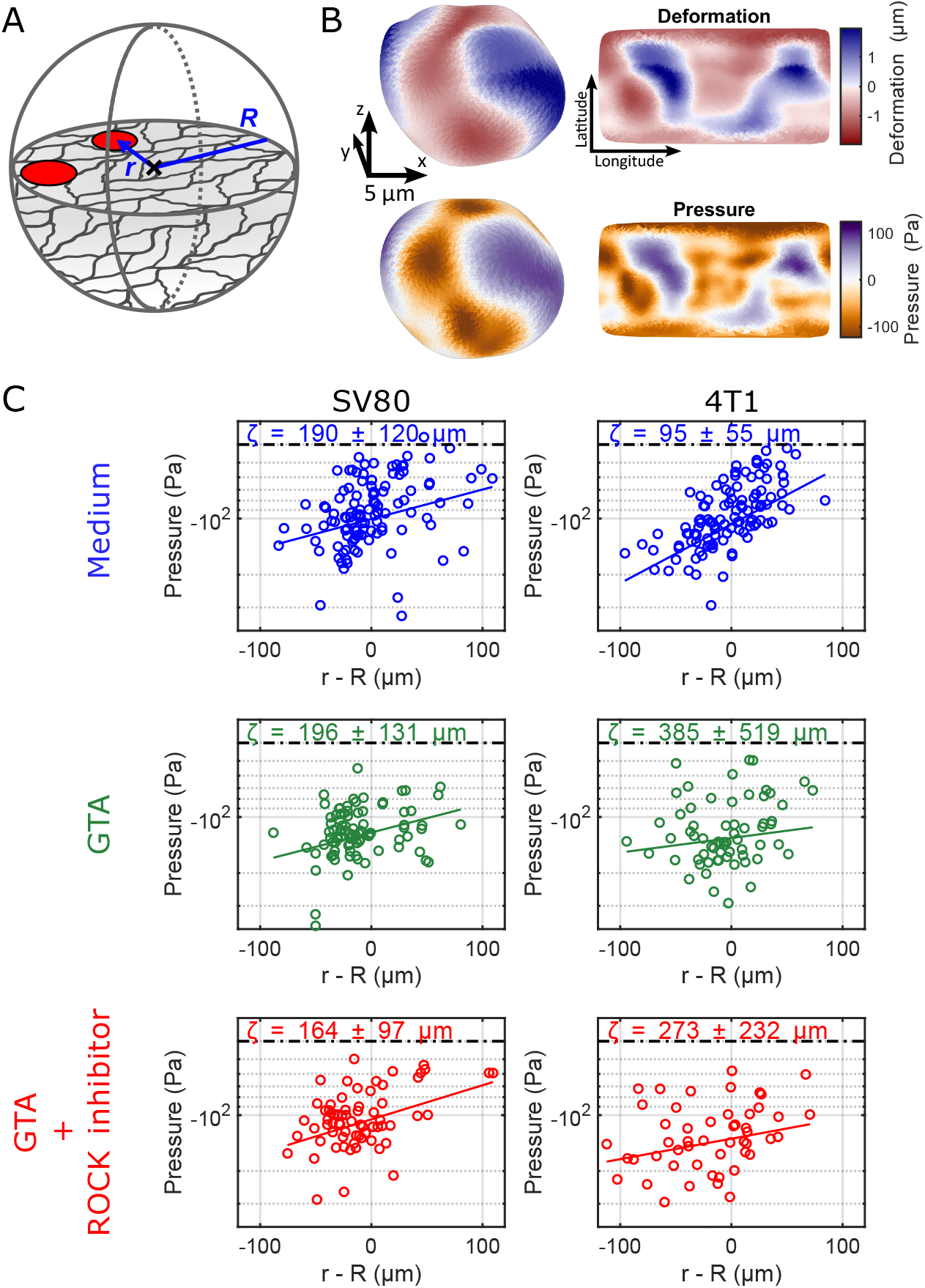
The internal spheroid pressure gradually decreased from its centroid and reached into the surrounding collagen matrix. **A**. Schematic overview of a spheroid of radius *R* with embedded microparticles (red) at a distance *r* from the centroid. **B**. Deformation (top) and pressure (bottom) field of a representative microparticle inside a spheroid. **C**. Mean pressure versus the distance to the spheroid edge (0; negative values inside the spheroid; positive values outside the spheroids in the ECM) for each of the experimental conditions. The solid line displays an exponential fit. A characteristic length scale *ζ* derived from this fit is displayed in each plot. The error shows the uncertainty of the fit. The dashed line shows the pressure offset (≈ *−*40 Pa) due to noise as measured for undeformed particles in the absence of spheroids.

To quantify a length scale up to which the spheroid-induced stress reaches into the ECM, we modeled an exponentially dropping absolute pressure as a function of distance *r* from the spheroid core,

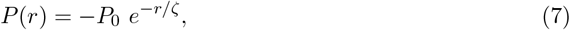

with core pressure *P*_0_, and characteristic length-scale *ζ* on which the spheroid-induced mechanical stress dissipates into the gel. In the < 10 Pa non-crosslinked gel this length scale was shorter for 4T1 than for the SV80 spheroids, indicating that the SV80 fibroblasts extended pressure further into the surrounding ECM. Upon cross-linking, the effective reach of the pressure was similar in the soft (“medium”, 190 ± 120 µm) and in stiff ECM (“GTA”, 200 ± 130 µm) for SV80 fibroblast spheroids. On the other hand, the pressure reach of 4T1 spheroids increased dramatically from 100 ± 60 µm in the soft ECM, to > 250 µm in the stiff, GTA cross-linked, ECM. Inhibition of ROCK had little effect on this behavior for SV80 and 4T1 spheroids.

These results showed that pressure levels measured using soft elastic hydrogel microparticles are significantly increased in the core of the spheroids. Mechanical stresses and pressures level off towards the outer spheroid layers and clearly extend beyond, reaching into the ECM. In line with the extensive ECM remodeling around spheroids we showed above, the extension of mechanical stress and pressure into the surrounding ECM reaches for hundred(s) of µm, lengthscales about or larger than the radius of a spheroid. While pressures caused by SV80 spheroids already penetrate into the ECM in soft environments, 4T1 spheroids react to ECM stiffening by an increase in their pressure reach.

## Discussion

Our study aimed to investigate how ECM stiffness affects stresses and pressures within tumors, how it affects cell migration and ECM remodeling, and how those mechanical and biological parameters are correlated. We show that pressure gradients within and around spheroids are dependent on cell type and, for tumor spheroids, also on ECM stiffness and ROCK activity. Both tumor and fibroblast spheroids respond similarly to ECM stiffness in terms of invasion, yet developing different migration patterns. Tumor spheroids exhibit a pronounced ability for collagen network remodeling, which decreases for stiffer ECM. In contrast, remodeling of the collagen network increases with ECM stiffness for fibroblast spheroids. Remodeling of the fibrous ECM is paralleled by build-up of stress and pressure gradients extending increasingly further into the ECM in stiffer environments. Molecularly, we show that ROCK inhibition reduces tumor spheroid invasion without affecting fibroblast migration or collagen remodeling.

In the three-dimensional context of tissue, it is not obvious how cell migration and tumor invasion depends on the mechanical properties of the surrounding ECM. Whereas an increased stiffness is known to trigger cellular programs that lead to higher cellular forces and mobility, simultaneously more stiff and dense environments will lead to a decrease in cell mobility. Indeed we here confirmed that the relation between stiffness, motility and invasion is more subtle. In our experiments we achieved stiffening of the low-density fibrous collagen matrix by chemical cross-linking. It should be mentioned that the mild cross-linking used, solely changed the elastic behavior of the gel. We observed no change in collagen fiber architecture, nor in fiber density (Fig. S4) in comparison to earlier reports suggesting that a cross-linked collagen matrix has a lower porosity than non-cross-linked controls [38]. Thus, any change in cell behavior could be attributed to mechanical properties of the environment. Cross-linking of the low-density collagen network led to a five-fold increase in storage modulus, consistent with earlier reports [39]. The increase in ECM stiffness elicited an increased invasion of tumor cells and of fibroblasts into the ECM. Those findings are in line with earlier related observations. It was reported that invadopodia formation and tumor-cell invasion was stronger in a stiffer matrix [15]. Likewise, interpolation of experimental results in 2D to the 3D situation in the current study indicates the stiffer ECM could provide a more rigid substrate for cancer cells to adhere to. Indeed such dependence was observed, in which cancer spheroids showed enhanced integrin adhesions in stiff ECM [16]. Cells likely transmit forces more effectively and develop stronger adhesion to the ECM when the ECM is stiff. Enhanced cellular integrin activity in stiffer ECM would enable the cells to resist detachment from the ECM and thereby promote their ability to invade. Interestingly, we observed a higher invasion potential for 4T1 spheroids as compared to SV80 spheroids. This observation is in line with the fact that 4T1 cells highly express E-cadherin, enabling them to develop stable cell-cell contacts. Those are the prerequisite to collective migration modes that we observed as finger-like structures in our confocal images. Collective cells have been reported to migrate faster than single cells [40].

The differential behavior between fibroblasts and 4T1 cells in terms of migration was reflected by the build-up of local stresses in and close to the spheroid, and the alignment of the collagen network around the spheroids. Local stress and pressure increased towards the spheroid core to about 100-200 Pa in the about 100 µm-radius spheroids investigated. There was no significant difference in core pressure for the different preparations. Even though the stiff matrix could potentially compress the spheroid and make the cells experience more stress, no increase in pressure was measured inside the spheroid.

It is obvious from the stress measurements as well as from the ECM remodeling, that the internal stresses significantly extend beyond the spheroid edge into the ECM in a smooth transition. The main difference between the different experimental conditions is how far the stress reaches out into the network. In the soft environment the reach of the mechanical stress of fibroblasts (≈ 200 µm) is significantly larger that that of the 4T1 cells (≈ 100 µm). Those stresses can lead to an alignment of the fibrous network of the ECM as observed in our experiments. In particular, 4T1 cancer spheroids show high, radial collagen alignment as well as rapid stress and pressure increase inside spheroids in soft matrices. High collagen remodeling was earlier attributed to local invasion [41], therefore it is surprising to see that the 4T1 spheroids invade less in matrices of high remodeling, compared to a matrix in which the collagen is less aligned. This suggests that a potential increase in ability to bind to the ECM rather than an increase in traction force generation needs to be considered in order to explain the reduced invasion capacity in low-stiffness gels.

Stiffening of the environment by chemical cross-linking results in an increase in mechanical reach of the stress beyond our observation volume. This suggests that the reach of traction forces is the result of increased stiffness and cross-linking of the matrix rather than the actin-myosin interactions of the cells themselves. In particular for the 4T1 cancer cells these results are not too surprising, as the efficiency of force transmission might be more relevant to invasion than the magnitude of these forces [42]. The fact that despite lower collagen alignment around the tumor spheroid, more migration is seen in stiffer ECM might be due to a regulatory mechanism to manage internal pressure within the spheroid. In contrast to cancer cells, non-cancerous fibroblasts typically show no invasion, have a normal metabolism, controlled cell growth and are responsible for producing extracellular matrix components like collagen and elastin [43].

The results above suggested that the force-generation machinery of individual cells would have only a minor impact on the stress distribution and remodeling capacity in the context of spheroids embedded in ECM. Inhibition of ROCK only slightly changed the mechanical reach and the ability for collagen fiber alignment. The reduction in collagen alignment and invasion observed for 4T1 spheroids in the presence of a ROCK inhibitor shows that these cells use ROCK to regulate their actin-myosin interactions. Indeed, it has been documented that ROCK promotes collagen remodeling and facilitates tumor cell invasion [44]. For SV80, however, we do not see a change in collagen alignment and invasion when adding a ROCK inhibitor, indicating that SV80 spheroids do not require ROCK for cellular force generation. Other kinases like MLCK can also be involved in regulating actin-myosin interactions and could therefore be used by cells instead of ROCK, which would explain opposite results to those shown by Beningo *et al*. [45].

The increased invasion of 4T1 tumor spheroids in the crosslinked matrix that is stiffer but does not show more prominent fibers suggests this is not related to more effective migration along aligned collagen fibers. Notably, our observations are made with spheroids embedded in a relatively soft 3D environment (< 100 Pa) gels. In a similar regime, a shift from 100 to 150 Pa promoted breast cancer invasion similar to our finding for 4T1 [5]. In much stiffer hydrogels (*≥* 2000 Pa) spheroid expansion is attenuated, and this is overcome by ROCK inhibition, [46]. In fact, a quantitative model showed that an intermediate ECM stiffness provides optimal conditions for cancer cells to polarize, contract and invade [47]. For cells attaching to a 2D substrate the stiffness regime is very different with stiffnesses of > 100 kPa promoting adhesion formation and migration [48]. This indicates that ECM stiffness and tumor invasion exhibit a non-linear behavior, potentially resembling a biphasic behavior with maximum invasion seen at medium stiffness. The behavior is influenced by the 2D verus 3D geometry and will likely be different for different cell types, which can explain the different response of 4T1 and SV80 spheroids to stiffening in our experiments.

## Funding

This work is part of the research program “The Active Matter Physics of Collective Metastasis” with project number Science-XL 2019.022, which is financed by the Dutch Research Council (NWO).

## Author Contributions

K.B., R.R.M., E.D. and T.S. designed the experiments. K.B. and R.R.M. conducted the experiments and performed data analysis. K.B., R.R.M., G.K., E.D. and T.S. wrote the manuscript. All authors reviewed and edited the manuscript.

## Disclosures

The authors declare no conflicts of interest.

## Supplemental Material

**Figure S1:**
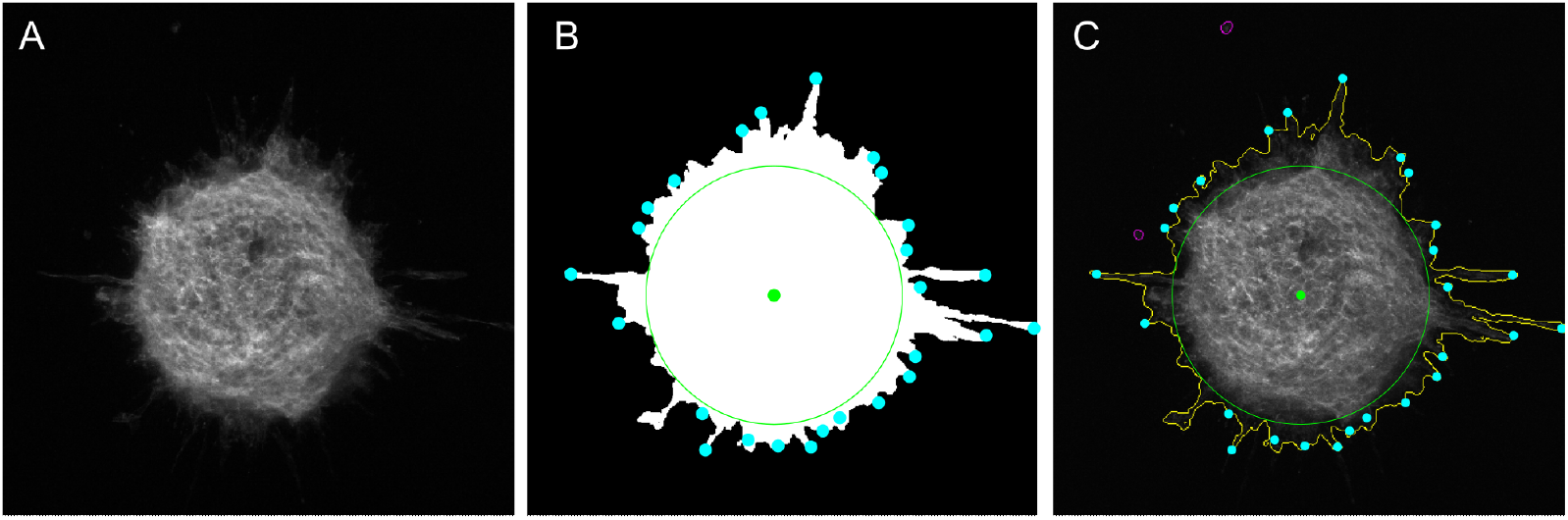
Analysis of 3D spheroid invasion assays by in-house Matlab migration app. **A**. A z-stack projection of a 4T1 spheroid was made by evaluating the variance of the signal in the z-direction. **B**. A mask of the spheroid including the endpoints of a Skeletonization shown in cyan. This mask was used to calculate the area, size of the core (green) and complexity. **C**. Overlay of the measure quantities with a maximum projection of the spheroid. The core radius is shown in green and the outline is shown in yellow. The branch area is the area of the spheroid outside the core.

**Figure S2:**
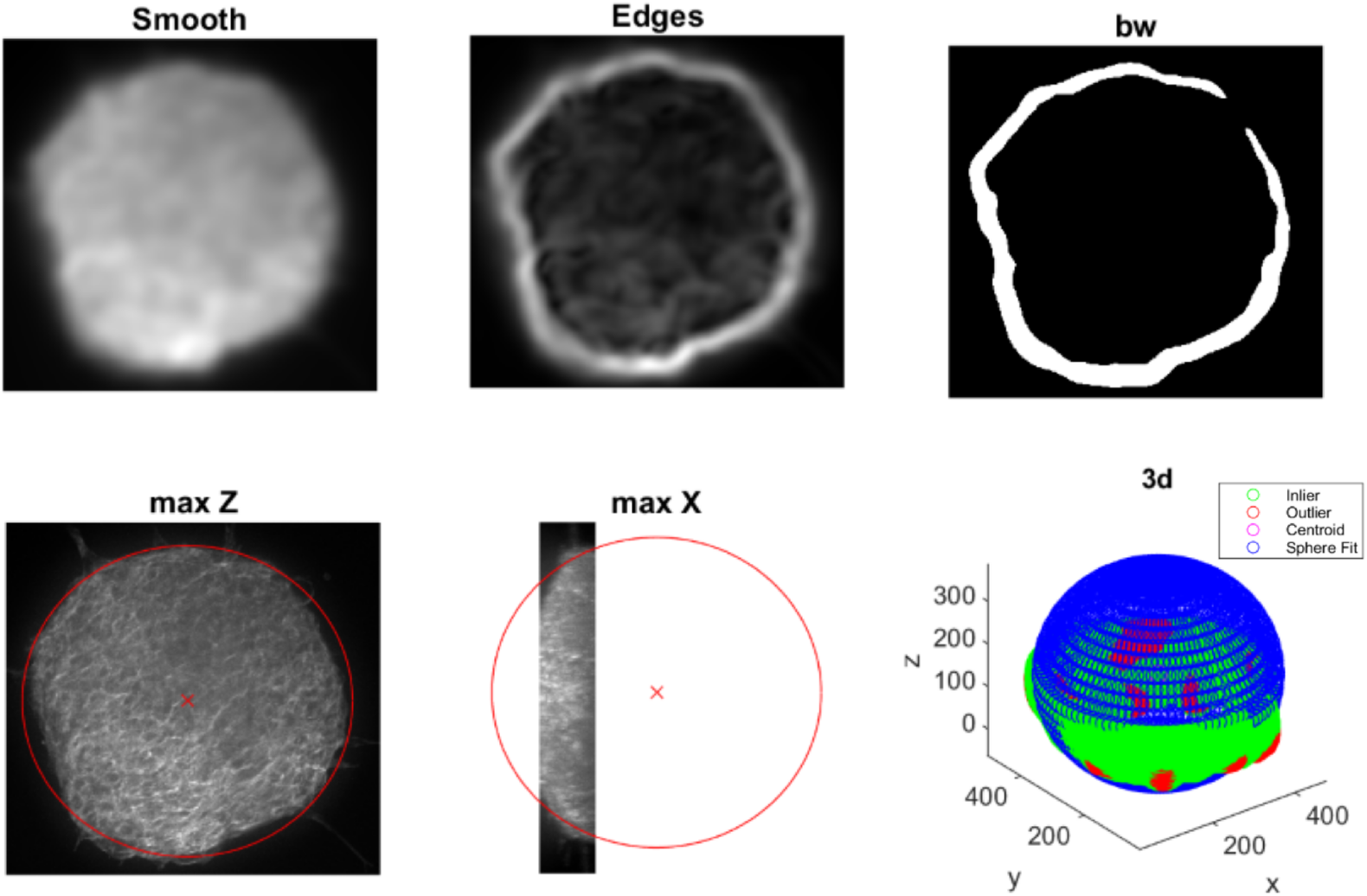
Spheroid center of mass (CoM) and boundary reconstruction. **A**. A Gaussian filter with a wide kernel was applied to the image stacks containing the actin cytoskeleton. **B**. A 3D Sobel operator was used to highlight the edges of the spheroid. **C**. The foreground was separated from the background to which a sphere was fit. **D-E**. Maximum z- and x-projection of the image stack. The resulting fit is displayed in red. **F**. 3D representation of the boundary fit (blue) to the edge-data (red and green).

**Figure S3:**
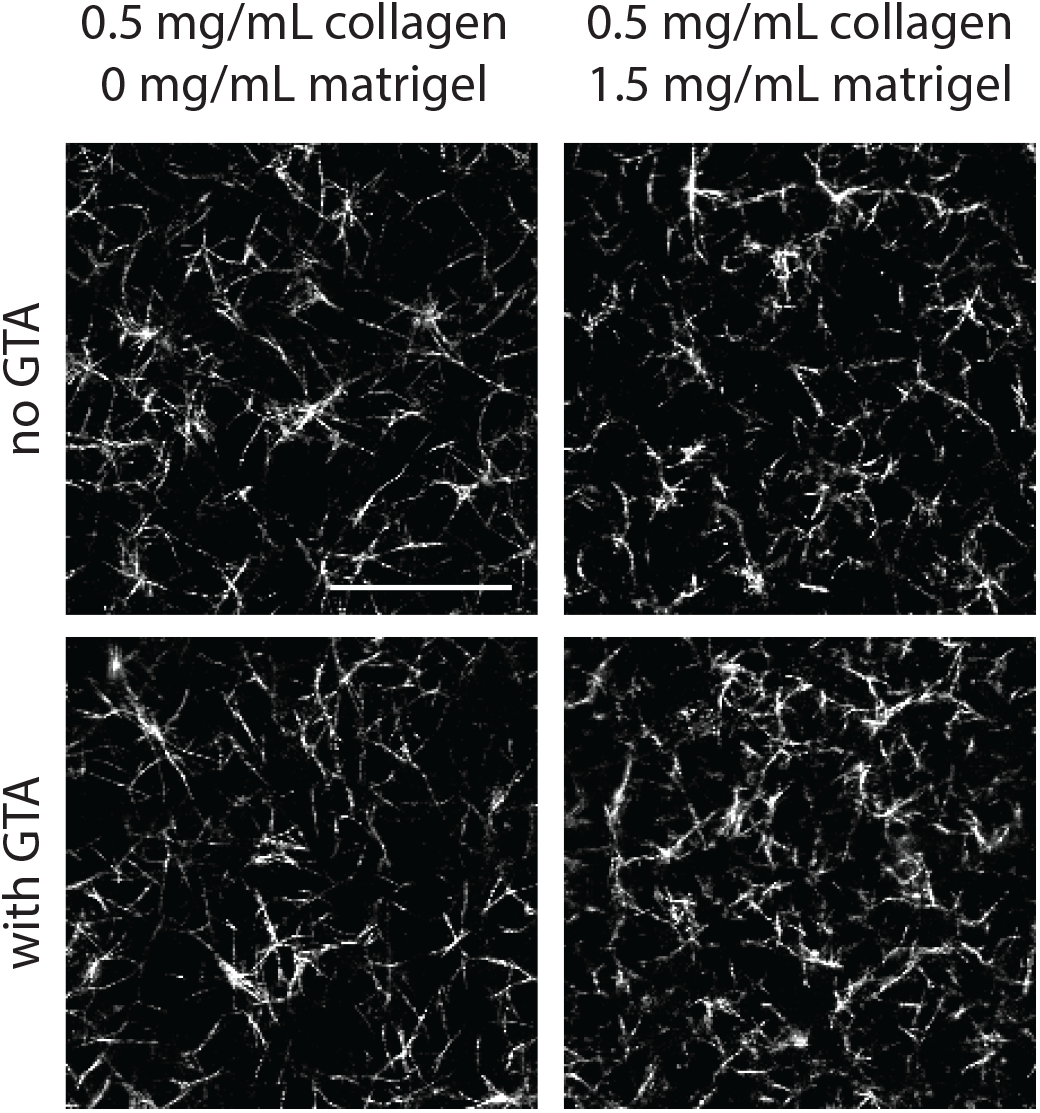
Reflection microscopy of collagen gels with and without matrigel. Collagen fibers (in white) have comparable lengths and thickness in gels with and without matrigel.

**Figure S4:**
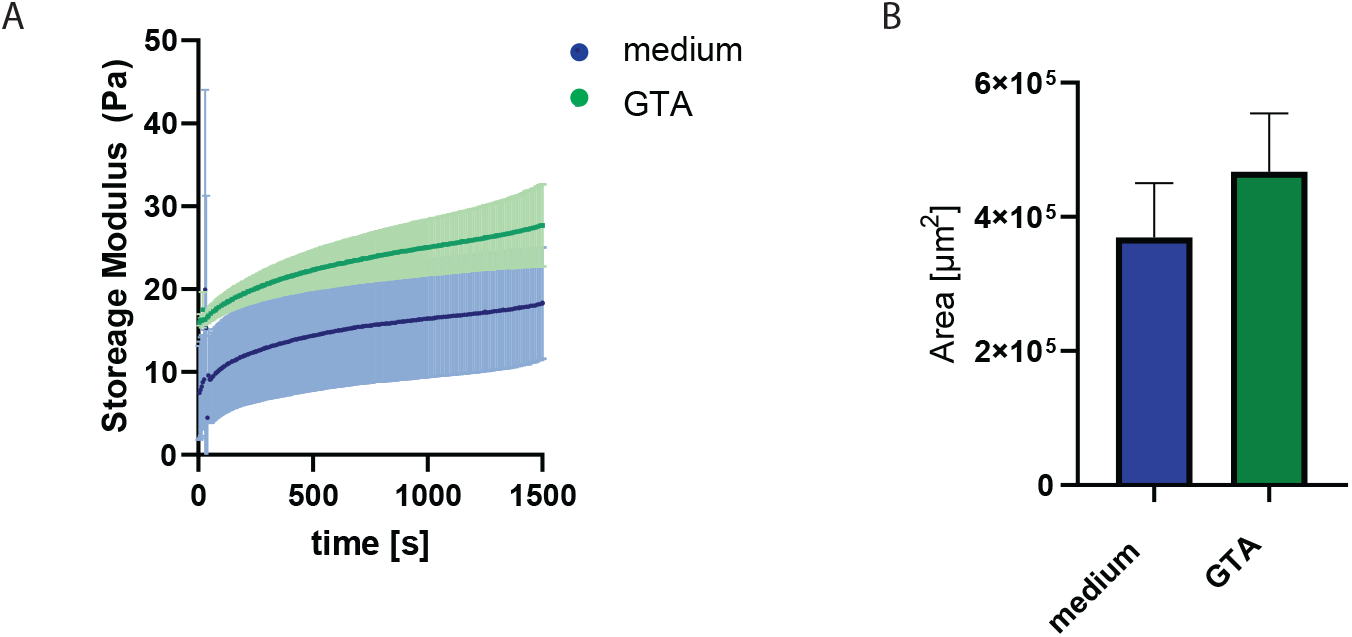
Rheology and fiber density of collagen gels. **A**. Storage modulus of gels before exposing to GTA, during collagen polymerization step. **B**. Quantification of area of reflection image covered by collagen fibers from three independent experiments.

**Figure S5:**
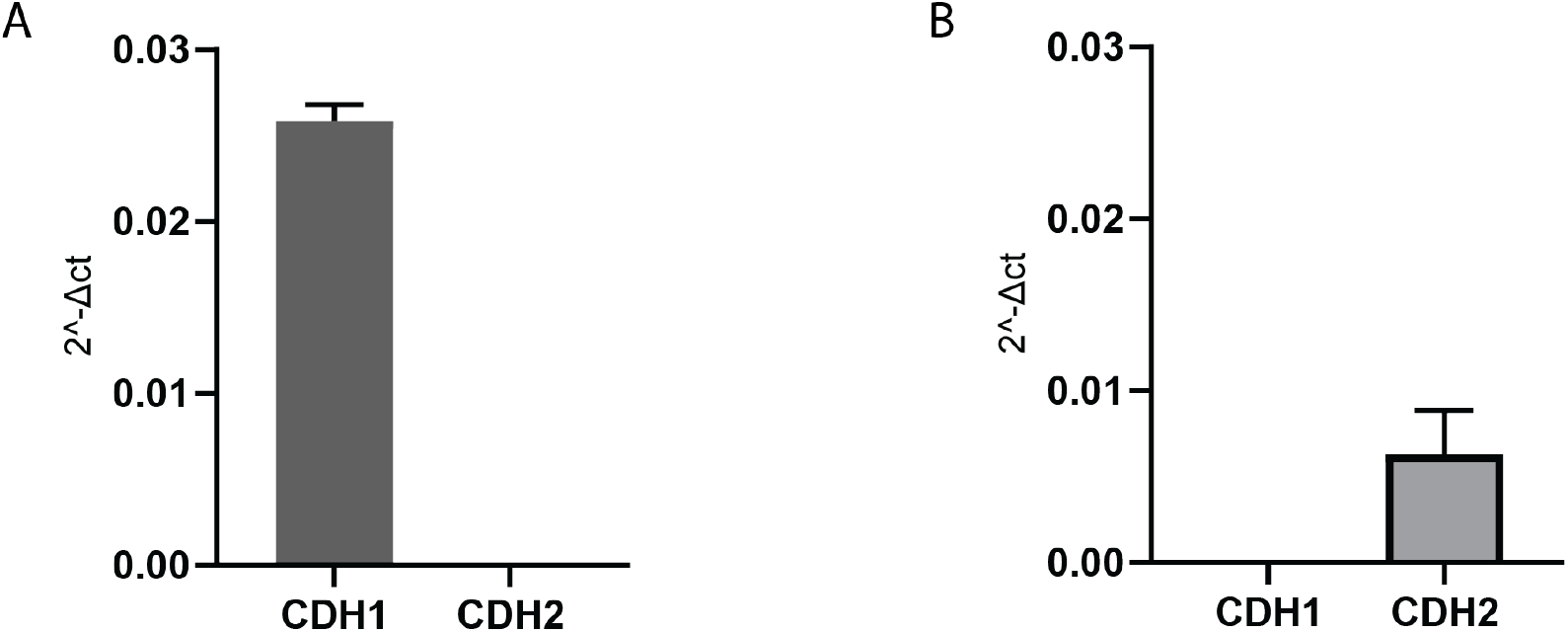
mRNA expression of Cadherins on SV80 and 4T1 cells cultured in 2D, normalized to Actin. **A**. 4T1 mRNA expression of CDH1 (E-cadherin) and CDH2 (N-cadherin). **B**. SV80 mRNA expression of CDH1 (E-cadherin) and CDH2 (N-cadherin).

**Figure S6:**
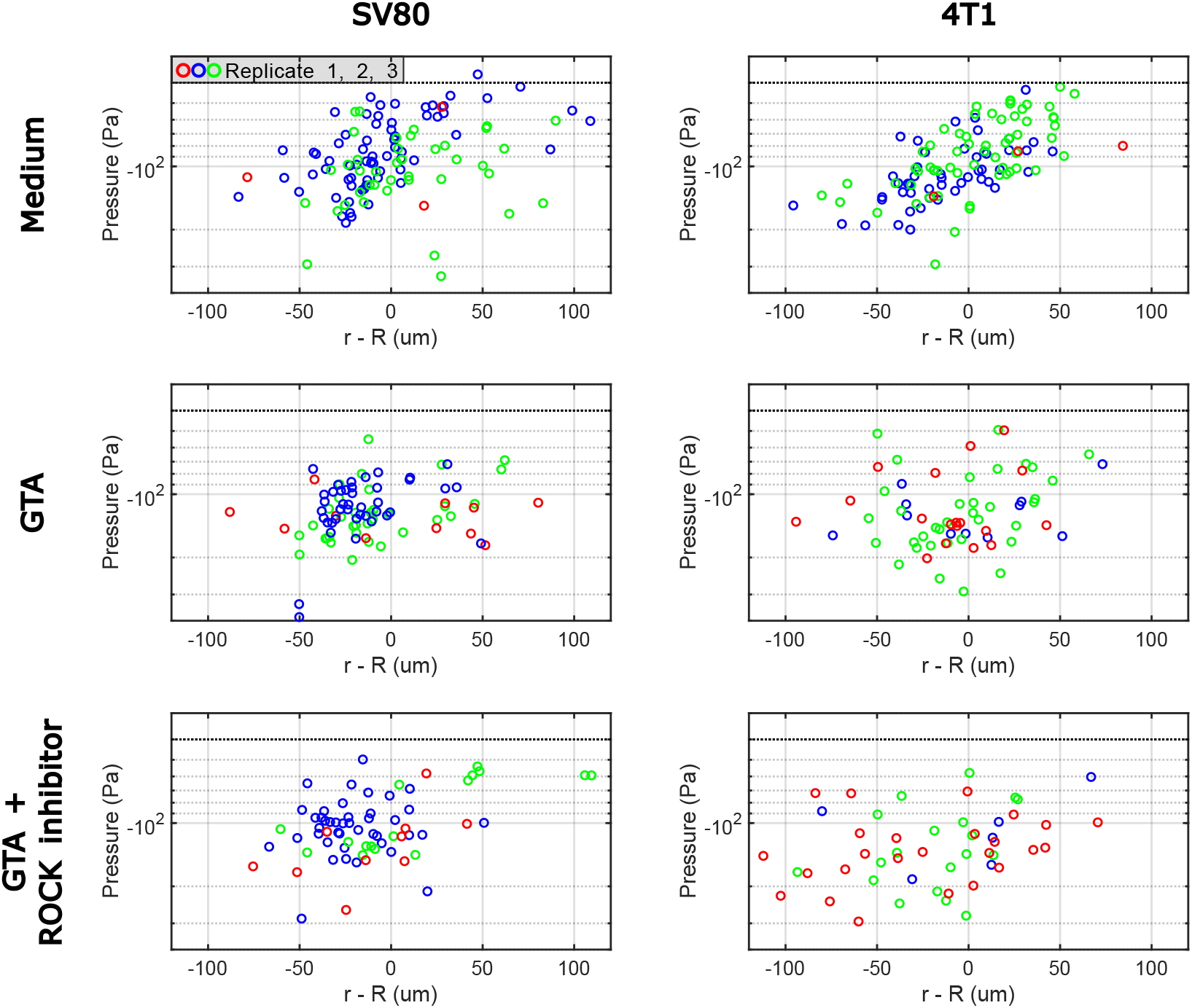
Separation of experimental replicates of Fig. 4 C.

## Notes

### Competing Interest Statement

The authors have declared no competing interest.

